## Supplementary material for "Trehalose supports the growth of *Aedes aegypti* cells and modifies gene expression and dengue virus replication": Table 1

Table 1. Trehalose-induced immunity genes

| Name | log2FoldChange | Description |
| --- | --- | --- |
| Cat | 1.171433 | Catalase |
| AAEL009467 | 1.738973 | coactosin-like protein |
| AAEL000488 | 1.7564 | interleukin-1 receptor accessory protein-like 1 |
| N/A | 1.960177 | defensin-A-like |
| PGRPLB | 1.844781 | peptidoglycan-recognition protein |
| GRRP  AAEL007374  AAEL009745  SRPN9  AAEL022496 | 1.274502  1.327801  1.85381  1.034501  1.013174 | holotricin-3  protein yellow  nitric oxide synthase  serine protease inhibitor  peptidoglycan-recognition protein |
